# Multi-omics analysis highlights the link of aging-related cognitive decline with systemic inflammation and alterations of tissue-maintenance

**DOI:** 10.1101/2025.07.13.662751

**Authors:** Stefano Flor, Thomas Dost, Madlen Haase, Rowena Simon, Simone Ederer, A. Samer Kadibalban, Jan Taubenheim, Maja Olecka, Alesia Walker, Johannes Zimmermann, Georgios Marinos, Sören Franzenburg, Philippe Schmitt-Kopplin, John Baines, Konstantin Riege, Steve Hoffmann, Lena Best, Christiane Frahm, Christoph Kaleta

**Author notes:** Shared senior authors (Lena Best, Christiane Frahm, Christoph Kaleta).

## Abstract

Aging-related cognitive decline is associated with changes across different tissues and the gut microbiome, including dysfunction of the gut-brain axis. However, only few studies have linked multi-organ alterations to cognitive decline during aging. Here we report a multi-omics analysis integrating metabolomics, transcriptomics, DNA methylation, and metagenomics data from hippocampus, liver, colon, and fecal samples of mice, correlated with cognitive performance in the Barnes Maze spatial learning task across different age groups. We identified 734 molecular features associated with cognitive rank within individual data layers, of which 227 features remain when integrating all data layers with each other. Among the single-layer predictors, several host and microbial features were highlighted, with host-associated markers being predominant. Host features associated with cognitive function mainly belong to innate and adaptive inflammatory activity (inflammaging) and developmental processes. Our findings suggest that cognitive decline in aging is tightly coupled to systemic, age-associated inflammation, potentially initiated by microbiome-driven gastrointestinal inflammatory activity, emphasizing a link between peripheral tissue alterations and brain function.

## Introduction

Cognitive decline in aging populations is a major public health concern, significantly contributing to increased morbidity and reduced quality of life in the elderly ^1^. While numerous factors are implicated in cognitive aging, the underlying biological processes driving neurodegeneration remain incompletely understood ^2^. Growing evidence points to a central role for the interplay between multiple physiological systems, suggesting that cognitive deterioration is not solely brain-intrinsic but arises from systemic dysregulation ^3,4^. Among these systemic factors, microbiota-derived metabolites have emerged as key mediators of age-related cognitive changes ^5^. Cross-species studies have revealed conserved gene regulatory changes across tissues during aging—such as enhanced inflammatory responses and suppressed cellular replication—suggesting that these processes may underlie both general aging and neurodegeneration ^6^.

In certain neurodegenerative diseases, such as Parkinson’s disease, it has been proposed that pathogenesis may vary between individuals, with some cases originating in the brain, and others in the gut (“gut-first” vs. “brain-first” hypothesis) ^7^. Several mechanisms have been suggested for microbiome-mediated neurodegeneration: microbial products that disrupt proteostasis and promote protein misfolding and aggregation ^8^; gut-to-brain translocation of misfolded proteins ^9^; and age-related gut barrier dysfunction leading to chronic, low-grade systemic inflammation ^10^. Bidirectional communication between the gut microbiota and the brain occurs through multiple pathways: direct neural signaling via the vagus nerve; modulation of immune responses and cytokine profiles; regulation of the hypothalamic–pituitary–adrenal (HPA) axis; and the production of neuroactive metabolites, including short-chain fatty acids and tryptophan derivatives ^11^. As demonstrated in a previous study, a hallmark of the aging gut microbiome in mice is a reduction in metabolic cooperation with the host ^12^. However, the significance of this reduction for aging-associated cognitive decline remains unclear, and few studies have systematically linked multi-omics, multi-tissue data to cognitive outcomes in aging.

In this cross-sectional study, we address this gap by using multi-omics data integration to investigate the role of multiple organ systems - including the gut and its microbiome - in cognitive decline during aging. Cognitive parameters were assessed in mice aged 3 to 28 months using the Barnes Maze for a spatial navigation task. Samples were collected for RNA sequencing (hippocampus, liver, colon); DNA methylation profiling (hippocampus and colon); shotgun metagenomics and untargeted metabolomics of fecal content. Constraint-based reconstruction and analysis (COBRA) modeling ^13^ and machine learning approaches were applied to identify factors associated with cognitive function. Changes in DNA methylation and gene expression, particularly those related to development and immune function, were revealed as the strongest predictors of cognitive decline. While the direct impact of bacterial metabolism on cognitive function appeared to be limited, our data showed a robust association between colon health and cognitive performance. These findings suggest that the gut microbiome may contribute to age-related cognitive decline indirectly—by shaping the local gut environment and modulating inflammation.

## Results

### Aging-associated cognitive decline in mice

We investigated the relationship between aging and cognitive function in a cohort of 83 C57BL/6J/Ukj male mice, comprising five age groups ranging from early adulthood to late age. To assess spatial learning and memory, we employed the Barnes Maze test (Figure 1A), a well-established task in which mice are trained to locate an escape hole on a circular platform using spatial cues. Two key behavioral measures were derived from this setup: primary latency, the time needed to locate the escape hole, and a cognitive score, a numerical representation of the search strategy used by the mice to reach the escape hole. These variables were recorded throughout six days of training (Figures 1B and 1C), during a probe trial (to assess short-term memory) where the escape box was removed, and during a retention test (to assess long-term memory) conducted several days later (Figure 1D). While the behavioral data from these cognitive tests in mice have been published previously ^14^, this study introduces a novel analytical framework that integrates these behavioral metrics into a single measure representing cognitive performance as a whole (Figure 1E). Following the approach in Ederer et al.^14^, we applied Tobit regression to the log-transformed latency data to account for right-censored trials, yielding model-based estimates of expected latency values. Similarly, we applied linear regression to the cognitive scores of the training period (Figure 1C). Unlike the previous approach, we retained both the slope and the intercept of the regressions as two separate cognitive features, indicating, respectively, initial performance and learning rate. In order to enable a direct comparison of the overall cognitive function across individuals, eight features per mouse were ranked within the cohort; the features included intercept and slope parameters from both primary latency and cognitive score regressions, as well as latency and cognitive scores from the probe trial and retention tests. The median of these ranks was used to compute a composite cognitive rank (Figure 1E; see Methods). This integrative approach enables a robust and nuanced comparison of cognitive function across individuals, revealing a clear decreasing trend over age (rho = -0.6664, p-val = 6.144×10⁻¹²), with significant differences emerging between younger (3–15 months) and older (24–28 months) groups (Figure 1E). This pattern is consistent with previous findings ^15^, that age-related cognitive impairment in C57BL/6J mice arises between 12 and 22 months of age, depending on whether the task assesses high or low attention levels, respectively. Mean-rank differences among age groups are listed in Supplementary Table 1.

**Figure 1:**
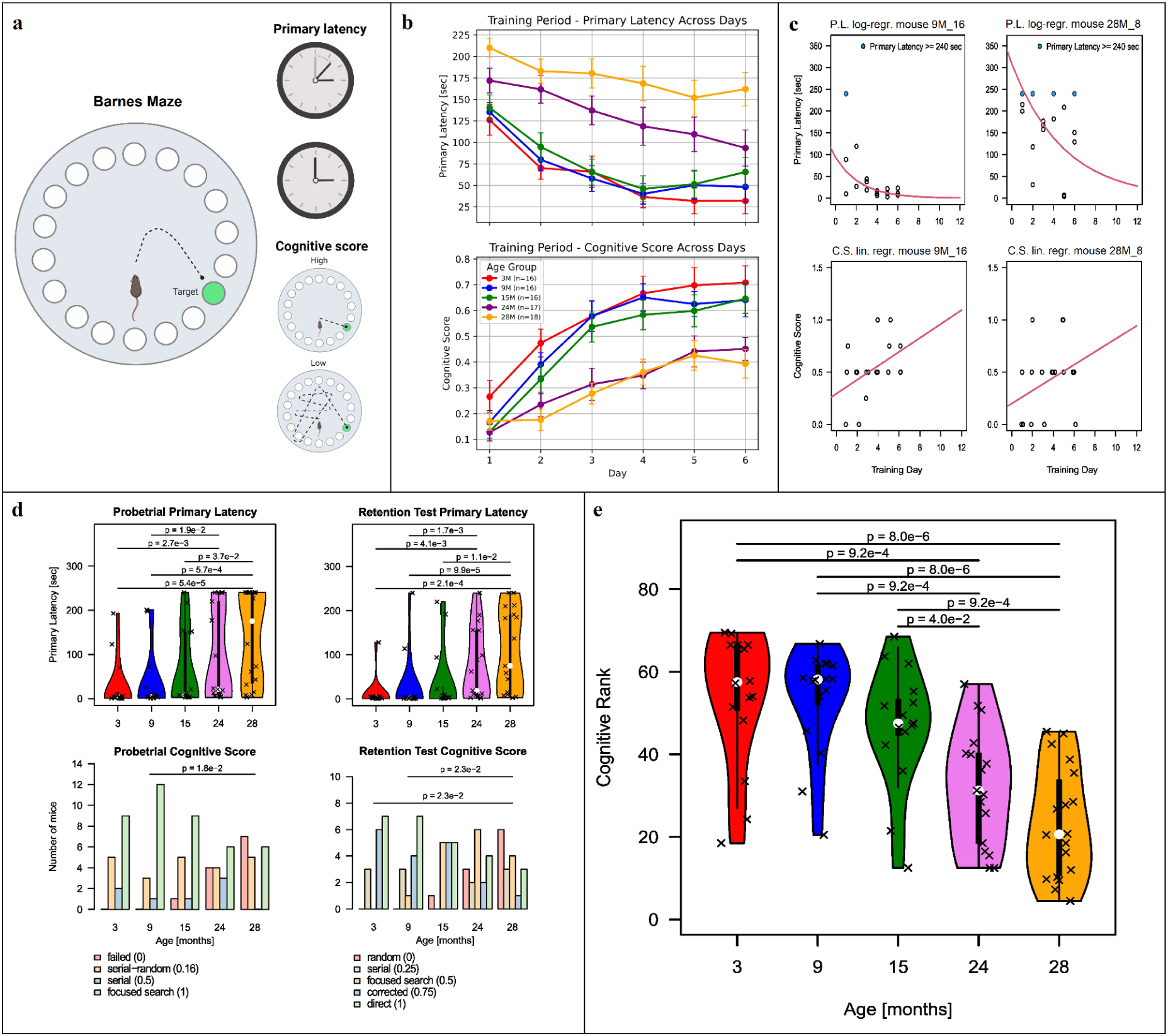
(A) Schematic representation of the Barnes maze setup, depicting primary latency (time that each mouse needed to locate the escape hole) and cognitive score (based on the search strategy that each mouse used to find the escape hole). (B) Line graphs of primary latency and cognitive score, stratified by age, across six training days. Adapted from Ederer et al.^14^. (C) Example scatter plots, showing logarithmic regression on the primary latency values and linear regression on the cognitive score across the training days for two mice, of age nine and 28 months. Blue dots indicate trials in which the mouse did not find the escape hole within the 240-second time limit. The training data has been collected for six days, the plot is extended to day twelve to illustrate the fitted models. (D) Violin plots and barplots of primary latency and cognitive score from the Probe Trial (short-term memory) and from the Retention Test (long-term memory). Statistical significance between groups is indicated.(E) Violin plots showing the distribution of the cognitive rank across five mouse age groups, with higher values representing better cognitive function. Dunn’s test p-values of the cognitive ranks’ distribution differences across age groups are displayed on top of the violins.

### Genome-scale metabolic modelling enables extensive characterization of gut microbiome’s functions

We reconstructed 383 metagenome-assembled genomes (MAGs) from shotgun sequencing data of fecal samples. Of those, 249 bins with a quality estimate of >=90% and a contamination estimate of <=5% were considered for further analysis to generate reliable metabolic models for downstream analysis. Most of the filtered MAGs belong to the phyla Bacillota (previously Firmicutes, n=159) and Bacteroidota (n=71). The MAGs from rarer phyla included Pseudomonadota, Campylobacterota, Cyanobacteriota, Desulfobacterota, Actinomycetota, Spirochaetota, and Verrucomicrobiota. Most of the discarded MAGs belong to the poorly characterized phylum Patescibacteria ^16^. To functionally annotate the assembled MAGs, we used gapseq ^17^ to reconstruct their corresponding genome-scale metabolic models. We obtained 249 metabolic models with an average of 1743 reactions and 1525 metabolites each, and a total of 4368 unique reactions and 3141 unique metabolites.

We quantified the metabolic potential of each mouse’s gut microbiome by generating a reaction abundance matrix, enabling functional annotation of microbiome activity. In this matrix, each column corresponds to a microbiome sample from an individual mouse, and each row reflects a specific metabolic reaction. Values indicate the relative abundance of reactions across microbiomes. This is based on diet-informed flux variability analysis simulations of each bacterium’s metabolic activity, integrated with MAG abundances. The final matrix included 2592 reactions, of which 166 are exchange reactions. Moreover, we predicted cross-feeding interactions in the microbiome of each mouse using the community flux balance analysis implemented in MicrobiomeGS2 ^18^, incorporating the relative abundances of each MAG from each fecal sample. For five community models, no feasible community flux balance analysis could be obtained. A total of 258 metabolites were predicted to have fluxes among the community members, between the community and the gut lumen, or both. The internal reaction fluxes were predicted to have a minimum of 759 reactions per community, a maximum of 1040, and a total of 3990 unique reactions over all communities in the cohort. We calculated the net flux for each exchange reaction within the community and between the community and the external environment, representing the gut lumen. Of 258 exchange reactions analyzed, 104 exchanges within the community and 63 exchanges with the lumen survived variance filtering (see Methods).

### Host tissue features are key predictors of cognitive rank

Host and microbiome features that are associated with the cognitive rank were selected with the Boruta ^19^ algorithm based on random forests (see Methods). We found 21 gene promoter methylation levels in the hippocampus and 98 in the colon to be related to the cognitive rank. Similarly, we identified 94 gene expression levels in the hippocampus, 93 in the colon, and 92 in the liver, as well as 20 MAG abundances, 27 reaction abundances, 5 exchange reactions within the bacterial community, and 15 metabolomics clusters that are predictive of the cognitive rank (Fig.2A). In order to select those features that are most predictive of the cognitive rank in the context of the overall multi-omics system, we used the features listed above in a second iteration of Boruta across all data layers simultaneously (see Methods). This resulted in the selection of 100 features from the gene promoter methylation data (2 from the hippocampus, and 98 from the colon) and 89 from the gene expression data (32 from the hippocampus, 34 from the colon, and 23 from the liver, as shown in Fig.2B). The following sections detail these features both within each individual data layer and in an integrative framework.

**Figure 2:**
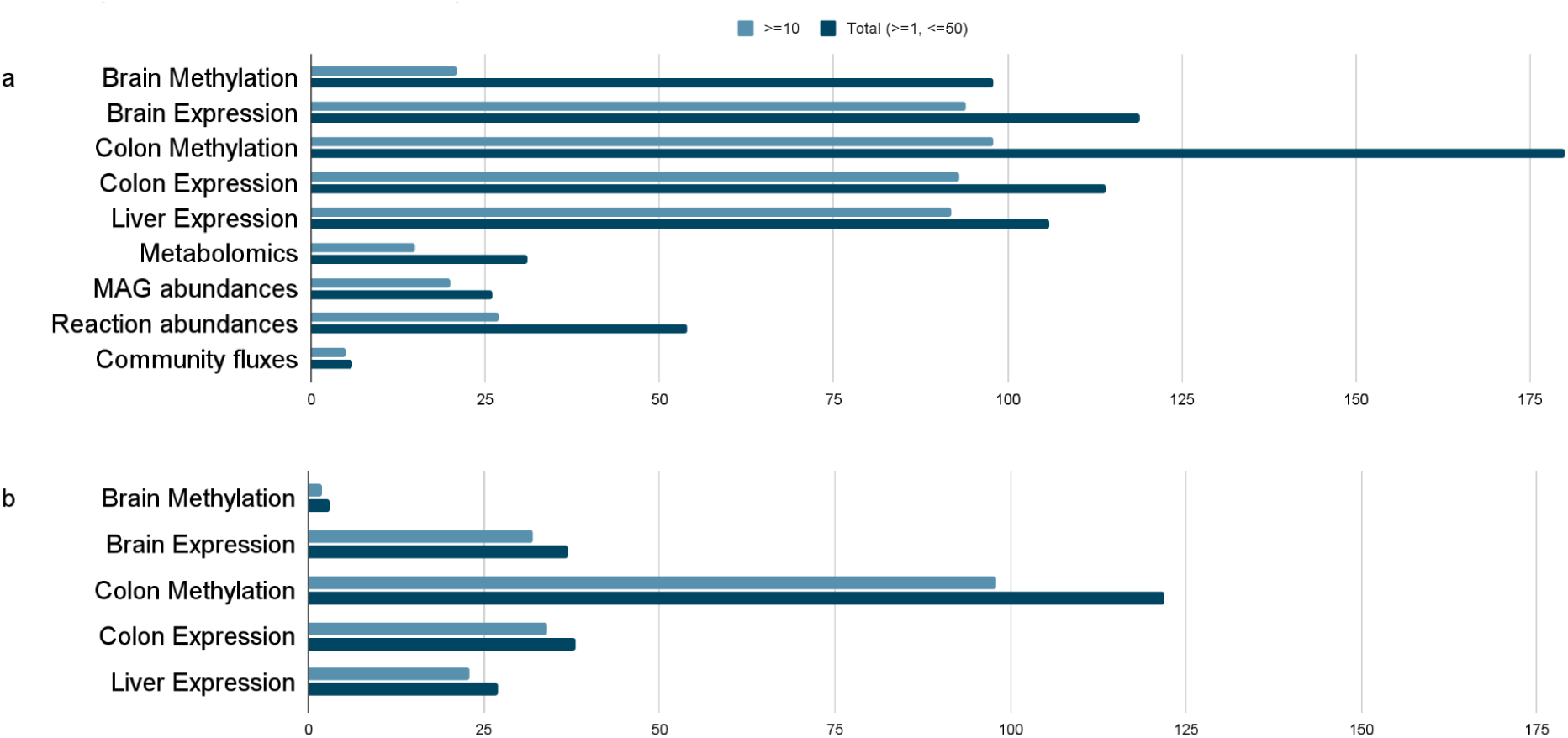
(A) Distribution of significant features identified by Boruta across individual data layers. Colors indicate the frequency at which features were selected in independent Boruta runs. (B) Features identified in the integrative analysis across all datasets. Colors indicate the frequency at which features were selected in independent Boruta runs.

### Host features predictive of the cognitive rank are involved in tissue development and immune function

The information on gene and promoter associations to functions and phenotypes that are presented in this section was collected through extensive cross-referencing of multiple databases (see Methods for the list of data sources) and scientific literature.

In the hippocampus, promoter methylation levels that are associated with cognitive function were primarily associated with genes enriched in developmental and regulatory processes (Fig. 3A). Of these, *Sox11*, *Lhx1*, and *Isl1* are transcription factors involved in morphogenesis and neuron differentiation. *Sox11* is essential for neurogenesis in mice ^20^ and has been associated with Coffin–Siris syndrome, a congenital disorder linked to mild intellectual disability in humans ^21^. Knockouts or mutations of *Lhx1* and *Isl1* have been linked to increased startle reflex and absent response to ear twitch, respectively.

**Figure 3:**
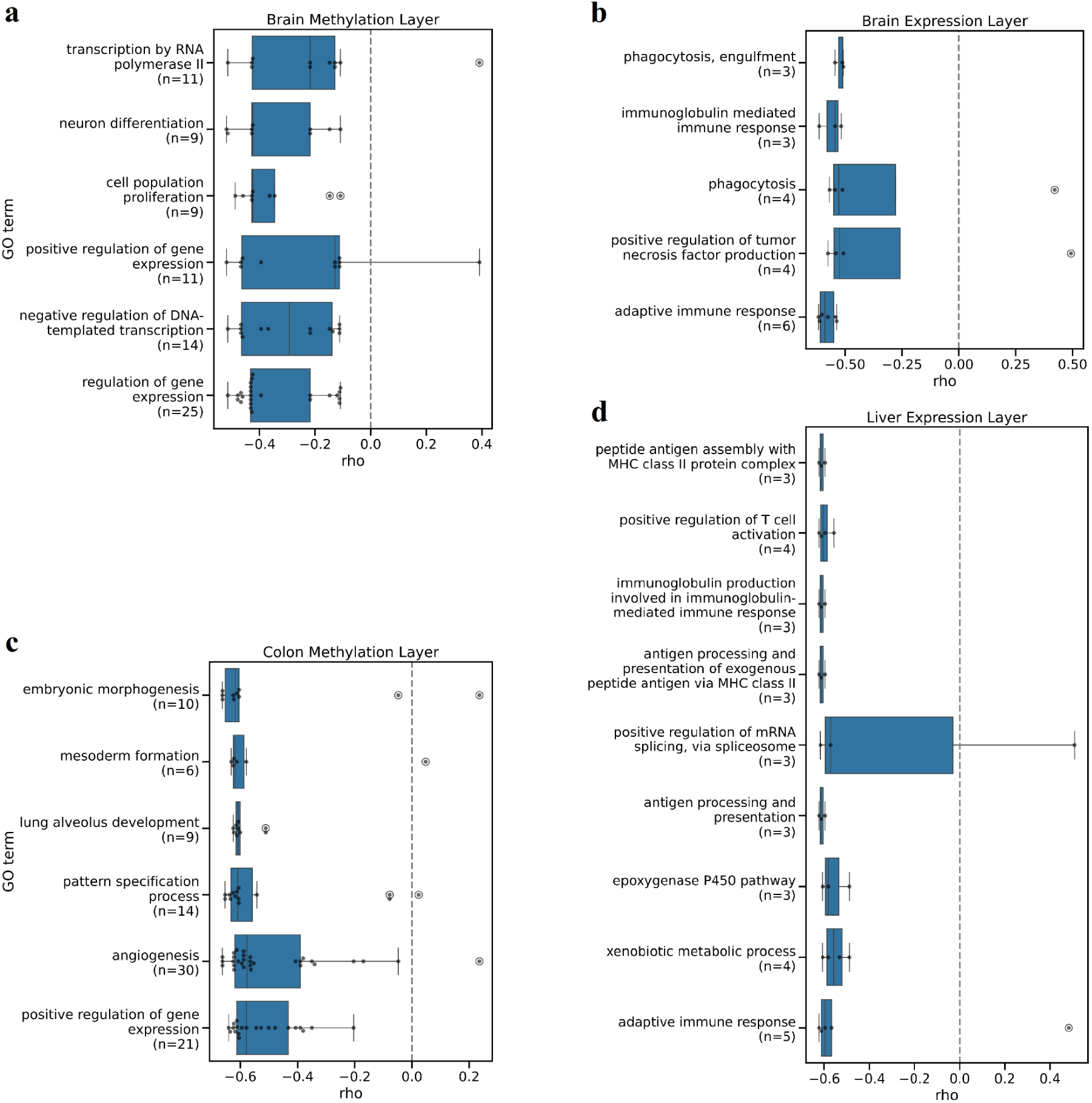
Gene Ontology enrichments in (A) brain and (C) colon methylation data layers indicate an inverse relationship between promoter methylation levels of developmental genes and cognitive rank. Dots in each box indicate the number of promoter regions of genes belonging to each GO term. Gene Ontology enrichments in (B) brain and (D) liver expression data layers indicate an inverse relationship between expression of immune genes and cognitive rank. Dots in each box indicate the number of transcripts of genes belonging to each GO term.

In the colon, genes whose promoter methylation was related to cognitive function were enriched for tissue development and were mostly negatively associated with cognitive rank (Fig. 3C). These included *Rora*, *Flt1*, and *Jam3*. *Rora* is known to drive gut inflammation by activating mTORC1 in T cells, making its high mucosal expression both a key pathogenic driver and a potential biomarker of anti-TNF treatment resistance in Irritable Bowel Disease in humans ^22^. *Flt1* (VEGFR-1) is primarily expressed in endothelial and immune cells (macrophages), where it mediates vascular and inflammatory responses ^23^. *Jam3* encodes a tight junction protein critical for gut tissue homeostasis ^24^, suggesting it may influence brain health by modulating gut permeability.

Hippocampal genes whose expression is related to cognitive rank are enriched for both adaptive and innate immune responses (Fig. 3B), consistent with inflammation being a key factor in neurodegeneration ^25^. Among cognition-associated genes, *Csf2rb*’s knockout has been linked to increased percentage center time in the open field test, and to an increase in eosinophils numbers, although only in female mice; while an increase in percentage center time can be interpreted as a decrease in anxiety, high eosinophils numbers are indicative of inflammation or infection. BRCA1 was also selected as related to the cognitive rank. In addition to its well-known role in breast cancer, this protein was shown to be essential for neuronal integrity and cognitive function; its depletion, triggered by amyloid-β accumulation, may contribute to cognitive decline ^26^.

In the liver, transcriptomic features related to cognitive function were enriched for immune processes, mRNA splicing via the spliceosome, monooxygenase activity, and xenobiotic metabolism (Fig. 3D). These included *H2-Aa*, *Lck*, *Ptk2b*, and *Skap1*, whose knockouts are associated with impairments in T-cells and macrophages function. Similarly, *H2-Ab1* knockout has been linked to immune dysfunction.

Gene ontology (GO) enrichment of cognition-associated colon transcriptomics features did not indicate any significant term. Nevertheless, we found several transcription factors implicated in inflammation and neurodevelopment among the selected features. Btn1a1 protein has been predicted to downregulate activated T cells as well as cytokine production. On the other hand, antibody cross-reactivity between Btn1a1 and Myelin Oligodendrocyte Glycoprotein is implicated in Multiple Sclerosis ^27,28^. Sept4 has been shown to attenuate alpha-synuclein neurotoxicity in the brain of a PD mouse model, and to be required for dopaminergic neurons to maintain key components of dopamine metabolism ^29^. Even if that has been shown only in the brain, considering Braak’s hypothesis ^30^, Sept4 may be relevant for a similar process in the colon, attenuating alpha-synuclein formation in the gut and its subsequent spread to the brain. Among the other transcription factors that we found among colon transcriptomics features *Irx2*, *Pou4f1*, *Pou6f1*, *Tfap2a, Ctdp1,* and *Msx1* are known to be involved in neuronal development or closely related processes ^31–37^, and *Fubp1* in brain cancer.

### Microbial features associated with cognitive function

To select microbial features that are significantly associated with the cognitive rank, we applied the Boruta algorithm (see Methods). Most MAGs whose abundance was related to cognition belong to the *Muribaculaceae*, *Lachnospiraceae*, and *Oscillospiraceae* bacterial families (Fig.4A, Fig.4B, Fig.4D). These species were predominantly positively associated with cognitive rank (Fig.4B, Fig.4D). It is worth noting that several members of the *Muribaculaceae* and *Lachnospiraceae* families are known to be generalist mucin monosaccharides consumers in gut communities, and to be protective against *C. difficile* infection ^38,39^. Among these MAGs, *CAG-485 sp002493045* from the *Muribaculaceae* family is the most abundant and most strongly associated with cognitive rank (Figure 4B). Recent research has demonstrated that metabolic interventions, such as empagliflozin (EMPA) treatment, can effectively restore levels of *CAG-485 sp002493045* in a mouse model of diet-induced obesity ^40^, making this bacterial species a potential target to be tested in the lab for microbiome-based interventions to maintain cognitive function in aging.

**Figure 4:**
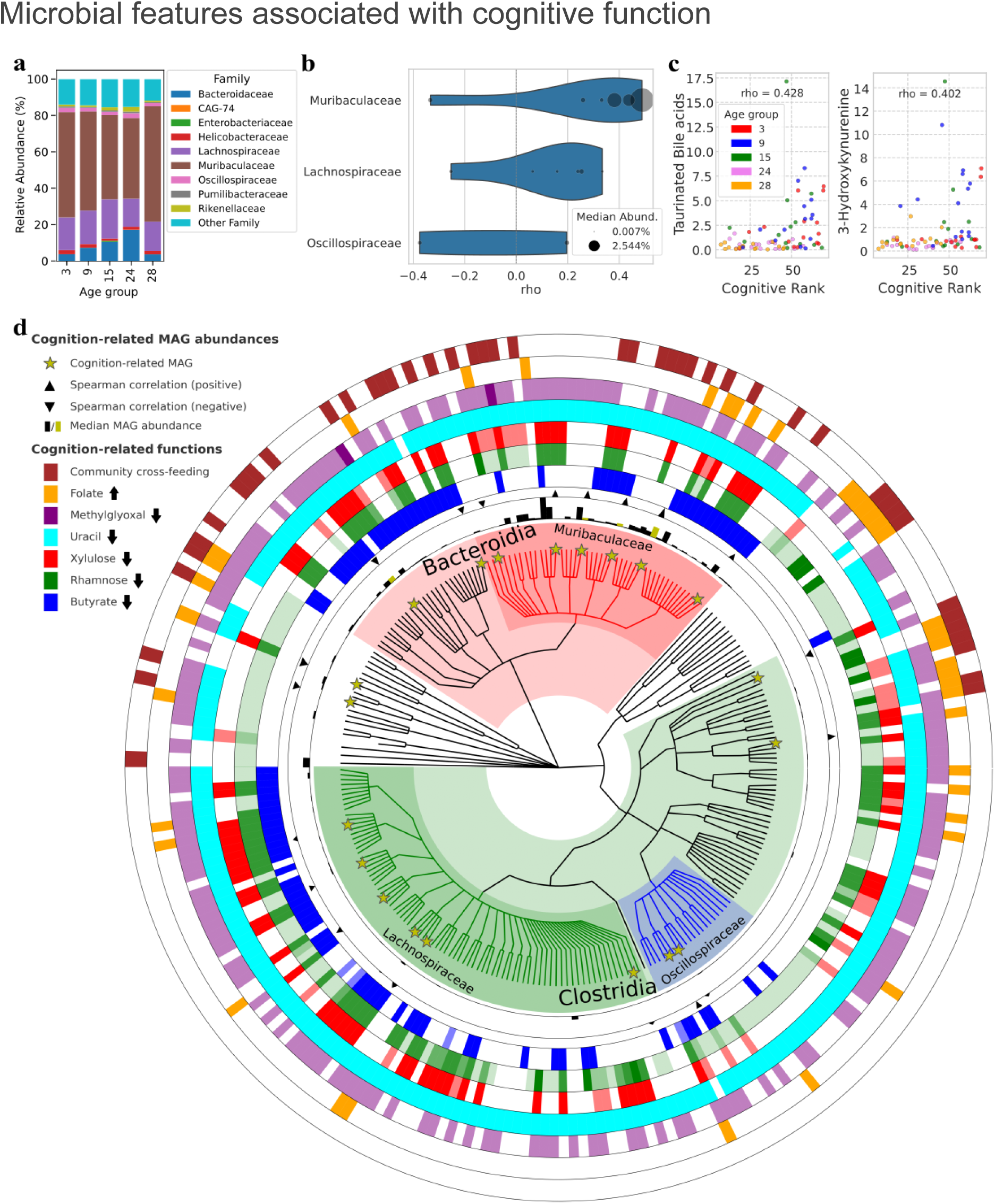
(A) Relative abundances of bacterial families of which at least one member is related to cognitive rank with remaining families grouped as ‘Other Family’. Samples are grouped by age. Six out of 52 MAGs belonging to *Muribaculaceae* have been selected by Boruta. The same applies to six out of 98 *Lachnospiraceae*, two out of 41 *Oscillospiraceae,* and one MAG for each remaining family. (B) Violin plot representing the distribution of Spearman rho values of the correlation between MAG abundances and cognitive rank for bacterial families with at least two Boruta-selected MAGs. (C) Boruta-selected metabolomics clusters levels (y axis) for each mouse, related to the cognitive rank and months of age. (D) Phylogenetic tree of MAGs selected by Boruta as predictive of cognitive function, generated with GraPhlAn and annotated with functional and abundance metadata. Branch colors indicate taxonomic affiliation, yellow stars mark MAGs selected by Boruta based on their relative abundance, and concentric rings show presence of metabolic functions in the gapseq models. Arrows in the legend indicate the direction of association of each annotated metabolic function with the cognitive rank. Ring color intensities reflect reaction counts, the black bars show median abundance with yellow-highlighted bars indicating those of Boruta-selected MAGs, and triangles indicate the direction of Spearman correlation between ten Boruta-selected MAG abundances and the cognitive rank: outward-pointing for positive correlations, inward-pointing for negative ones.

After identifying MAG abundance profiles related to cognitive function, we applied two complementary approaches to infer active metabolic reactions within the microbiome. First, we inferred abundances of metabolic reactions for each mouse’s microbiome community (see methods). We identified 27 reactions whose abundance is predictive of cognitive rank. The full list of reactions and their association with the cognitive rank is displayed in Supplementary Table 2. Reactions related to butyrate, rhamnose, propanediol, xylulose, uracil, and 2-oxopropanal (methylglyoxal) metabolism are inversely correlated with cognitive rank. The negative association of butyrate-producing reactions and spatial learning performance observed in our aging mouse cohort aligns with findings by Sampson et al. ^41^, which indicates that SCFAs (short-chain fatty acids) exacerbate neuroinflammation and α-synuclein aggregation in a Parkinson’s disease mouse model. Interestingly, propanediol exposure during development has been shown to cause negative behavioral effects in a zebrafish cohort ^42^. Rhamnose utilization can further increase propanediol levels, as the latter is an intermediate of propionate production from rhamnose through the propanediol pathway in gut bacteria ^43^. Methylglyoxal has been associated with lower grey matter volume, but not with white matter or hippocampal volumes, in an elderly human cohort ^44^. The only reaction whose abundance shows a positive association with cognitive rank is folate transport, aligning with previous findings demonstrating that the gut microbiome can modulate folate availability in Parkinson’s disease ^45^.

In a second approach, we constructed a community model for each mouse microbiome, enabling us to identify reaction activities for each microbe in the community, as well as exchange fluxes within the microbial community and with the external environment (i.e. the host’s colonic epithelium). Cross-feeding of a few molecules —including heme, S-adenosyl-homocysteine, S-adenosyl-methionine, and D-ribose—between bacteria were selected as important features, whereas no exchange flux with the gut lumen was selected. Most of these reactions are found in *Bacteroidia* (Fig.4D); this pattern suggests that members of this class may play a central role in mediating community-level metabolic interactions associated with cognitive function.

We found 15 fecal metabolomic clusters to be associated with cognitive rank, of which two could be identified as taurinated bile acids and 3-hydroxykynurenine; both of them showed a positive association with cognitive rank (Fig. 4C). Although 3-hydroxykynurenine (3-HK) is generally known for being part of the neurotoxic branch of the kynurenine pathway, it has been shown to be related to lower odds of cognitive impairment in pre-diabetic and type 2 diabetic patients ^46^.

### Multi-Omics Signatures of Cognitive Decline

Integrating the single-layer hits, our feature selection procedure identified a refined set of 227 host features (Figure 2B); of these features, 189 were selected ten or more times across the 50 feature selection runs (detailed feature selection procedure in Methods), and are displayed in Figures 5 and 6. Data layers related to the host tissues are the ones that carry relevant information when all omics layers are taken into consideration simultaneously, as no features from metabolomics, bacterial, or modeling data were selected in this run (Figure 2B). In the single-layer analysis (Figure 2A), colon methylation features were selected at similar rates as features from colon, liver, and brain expression layers. In contrast, the multi-layer analysis revealed a more than twofold increase in selected colon methylation features, highlighting the strong relevance of epigenetic changes in the colon for cognitive aging.

**Figure 5:**
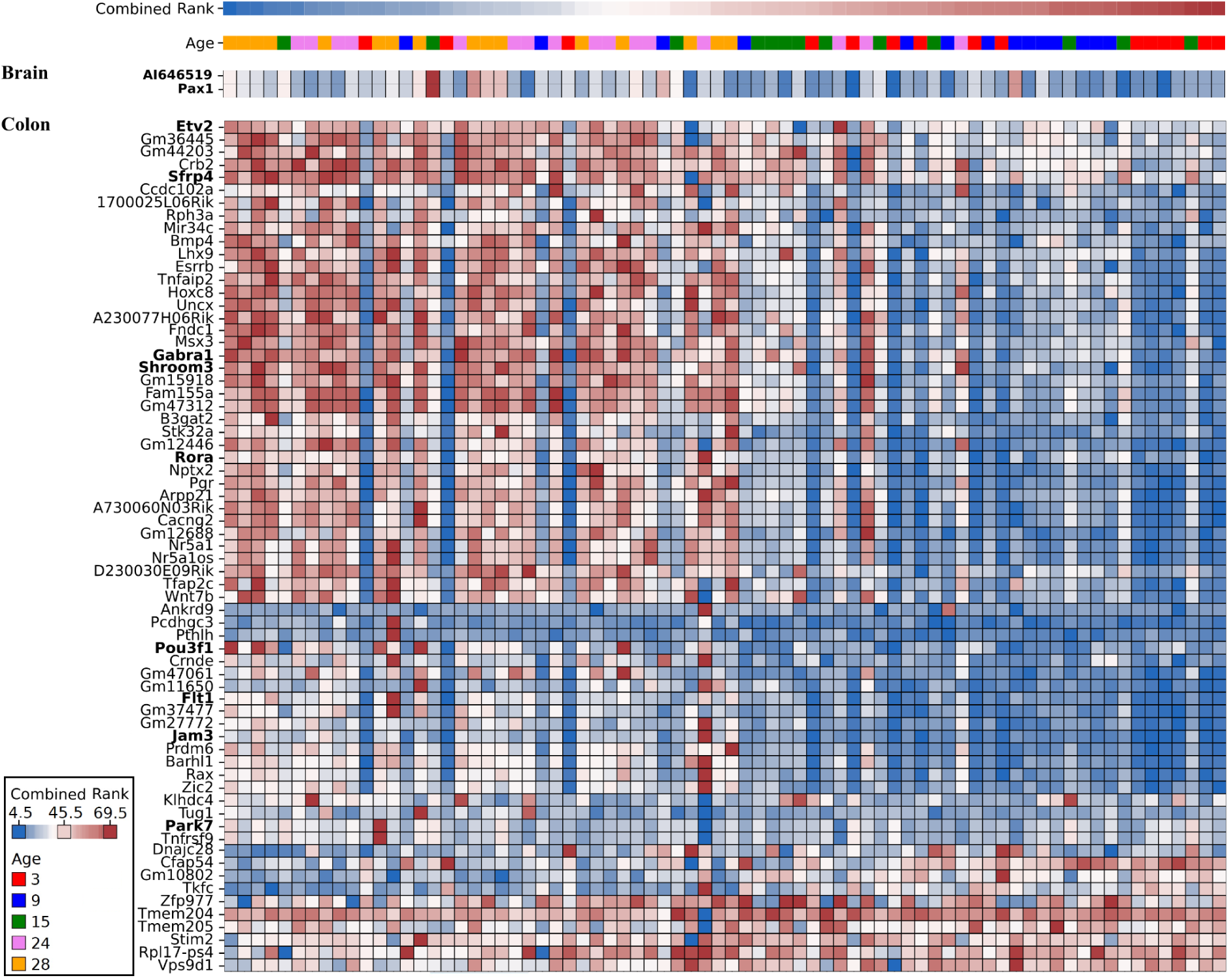
Brain and colon gene promoter methylation levels selected by Boruta ten or more times in the combined layers analysis. Each column represents a mouse. Combined cognitive rank and age information are layered on top of the gene promoter methylation heatmap, their values are shown in the legend at the bottom-left of the figure. Methylation levels are min-max normalized row-wise (gene-wise). Dark blue, white and dark red indicate the lowest, mean and highest expression values respectively. Genes discussed in this paper for being known to be involved in cognitive function, behavior, immunity, or developmental processes are indicated in bold

**Figure 6:**
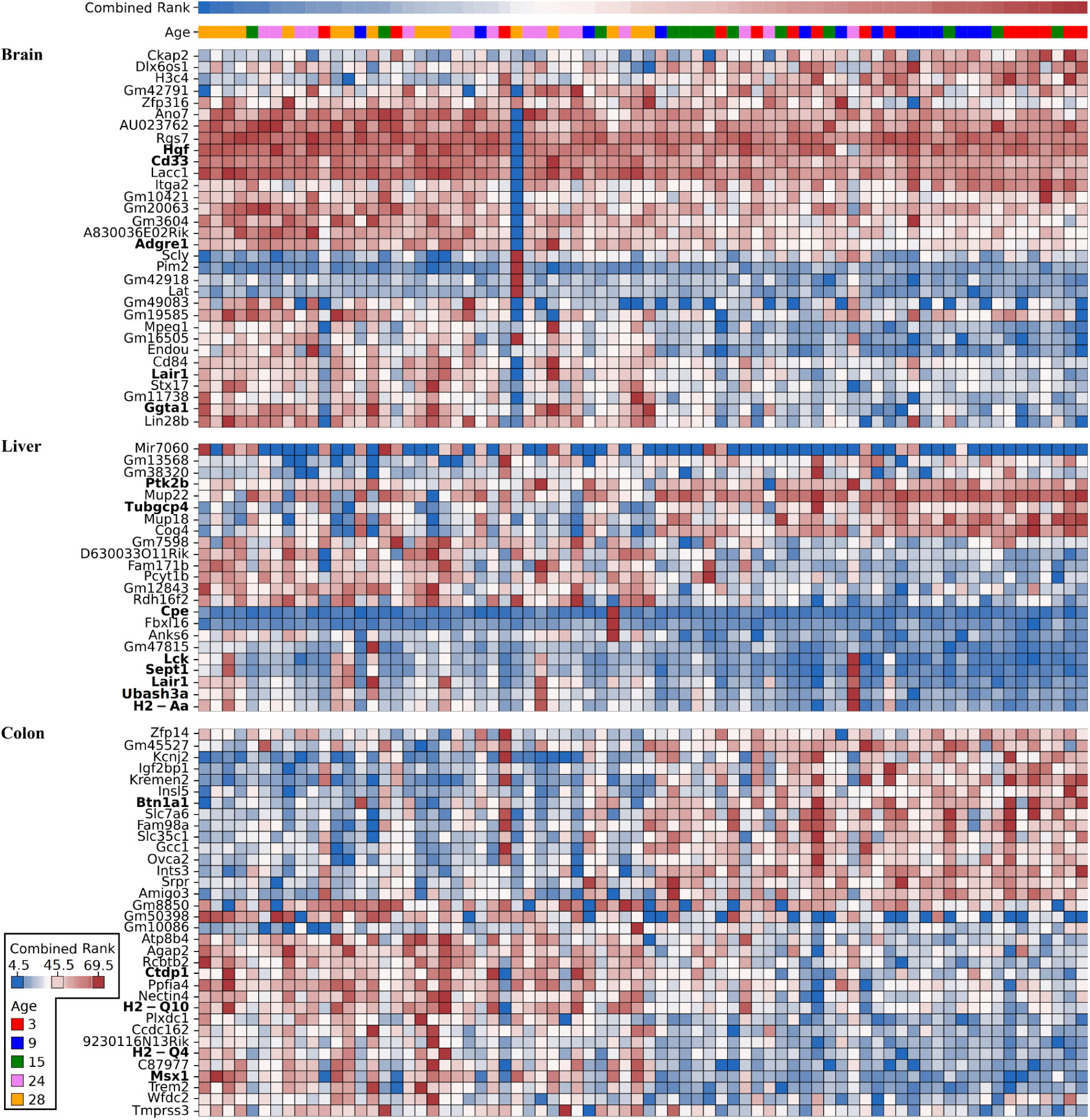
Brain, colon and liver gene expression levels selected by Boruta ten or more times in the combined layers analysis. Each column represents a mouse. Combined cognitive rank and age information are layered on top of the gene expression heatmap, their values are shown in the legend at the bottom-left of the figure. Expression levels are min-max normalized row-wise (gene-wise). Dark blue, white and dark red indicate the lowest, mean and highest expression values respectively. Genes discussed in this paper for being known to be involved in cognitive function, behavior, immunity, or developmental processes are indicated in bold

Methylation levels of the genes *Flt1*, *Rora*, and *Jam3* in the colon, which were already described in the single layers section of the results, were selected again in this run. Other genes previously implicated in neural and vascular development—*Shroom3*, *Etv2*, and *Pou3f1*—were also identified among the selected features. *Shroom3* plays a critical role in neural tube morphogenesis, with its disruption leading to defects in brain organoid models and linking it to neurodevelopmental vulnerability ^47^. *Etv2* acts as a pioneer transcription factor that remodels chromatin to drive endothelial specification by recruiting BRG1 to otherwise inaccessible genomic regions ^48^. *Pou3f1* facilitates neural differentiation by activating neural lineage programs and suppressing signals that inhibit neural fate ^49^. Notably, *Park7* methylation emerged as a predictive feature from the integrated analysis, though it was not highlighted in the single-layer analysis because it was selected in fewer than ten runs of the feature-selection algorithm (see Methods). *Park7* is involved in idiopathic Parkinson’s disease, maintaining gut microbiome balance, and regulating intestinal inflammation, making it a potential therapeutic target for gut-brain-axis disorders ^50–52^. Activation of epithelial gamma-aminobutyric acid (GABA) A receptors α1 (encoded by *Gabra1*) worsens DSS colitis by inhibiting mucosal regeneration ^53^. In contrast, promoter hypermethylation of *Sfrp4* distinguishes ulcerative-colitis–associated dysplasia/cancer from non-inflamed mucosa, reflecting chronic Wnt-driven inflammatory injury ^54^. As for the brain methylation data layer, only *Pax1*, which is a developmental gene which drives spine formation during fetal development ^55,56^, and *AI646519*, a lncRNA expressed on the reverse orientation of *Pax1* promoter, were selected as predictors of cognitive rank.

In total, 32 brain, 23 liver, and 34 colon expression features were selected (Figs. 2B and 6). Among genes expressed in the hippocampus, *Lair1*, *Ggta1*, *Hgf, Cd33 and Adgre1* have known roles in immune function and brain stress responses. *Lair1*, whose expression is negatively correlated with cognitive rank, regulates immune responses by inhibiting immune activity against self-recognized cells and interacts with collagen ^57^. This link to collagen —which undergoes structural and functional deterioration with age due to enzymatic cleavage, accumulated cross-links, and complement protein binding—is being investigated as a therapeutic target in neurodegenerative diseases ^58^. *Ggta1* responds to cerebral ischemia and its downregulation has been linked to protective preconditioning effects ^59^. *Hgf* is linked to brain development ^60^ and elevated tau biomarkers relevant to Alzheimer’s disease ^61^, and is known to be involved in repair of several tissues and organs including the brain ^62^. CD33 modulates the immune response in microglia and influences amyloid-beta clearance ^63^ while *Adgre1* is a microglial inflammation marker ^14^. In the liver, *Tubgcp4* is one of the few genes that positively correlates with cognitive rank, and is linked to brain and eye development disorders, such as microcephaly and chorioretinopathy ^64^. Another gene positively associated with cognitive rank is *Ptk2b*, encoding the neuronal kinase PYK2. This gene has been linked to amyloid-beta accumulation and cognitive impairment in the brain ^65^, though its liver expression may reflect systemic effects. Other notable liver genes include *Ubash3a*, a regulator of T-cell function linked to autoimmunity ^66^ that also has a role in tissue development ^67^ and *Cpe*, which is known to synthesize a neurotrophic factor involved in cognitive resilience against stress and has been shown to prevent neurodegeneration and memory loss in Alzheimer’s mouse models ^68^. Reduced quantities of synaptic vesicles in the retina have also been reported in *Cpe*-mutant mice, further linking this gene to neurodegeneration ^69^. Similarly to *Tubgcp4* and *Ptkb2*, it is not trivial to infer how expression of these genes in the liver could affect cognitive function, but may be indicative of a systemic alteration in the expression levels of developmental genes. *Sept1* maintains Golgi integrity in hepatocytes, supporting liver development ^70^, while elevated expression of *Lair1* on circulating monocytes marks early cirrhosis ^71^.

Colonic expression levels of *Btn1a1*, *Msx1*, and *Ctdp1*, previously described in the single data layer feature selection, were reaffirmed in this integrated analysis, together with *H2-Q10* and *H2-Q4*. The latter are part of the non-classical MHC class Ib family, involved in immune regulation. *H2-Q10* has been implicated in the development of liver-resident natural killer and gamma-delta T cells ^72,73^, while *H2-Q4* is involved in antigen presentation to the immune system, and has been shown to take part in rapid immune responses following lipopolysaccharide (LPS) exposure in an astrocyte subtype of the mouse brain ^74^. Methylation and expression levels of the genes that we mentioned here or in the previous sections are highlighted in bold in figures 5 and 6.

## Discussion

In this study, we investigated the biological processes underlying age-related cognitive decline across multiple omics layers in a mouse cohort that spanned from very young to very old individuals. To identify all relevant features across our high-dimensional multi-omics and multi-tissue dataset, we performed feature selection with random forests implemented in Boruta ^19^.

Several of the genes that we found to be involved in cognitive decline have been previously related to cognitive and behavioral alterations shown to be implicated in neurodevelopment and immune function. Abnormal DNA methylation, other epigenetic changes, and mutations in neurodevelopmental genes are known to contribute to neurodegenerative diseases and aging-related cognitive decline ^75–79^. Developmental genes are not only crucial for cell differentiation in the developing organism, but also for long-term maintenance of homeostasis as well as for tissue repair during wound healing and aging ^78–80^. In highly regenerative species such as the axolotl, reactivation of developmental programmes drives scar-free regeneration ^81^, whereas in mammals, the response is more limited. Mouse studies summarised by Eming et al. ^82^ show that cutaneous and sterile liver injury proceed through an early pro-inflammatory phase, followed by a reparative phase in which macrophages adopt anti-inflammatory, growth-promoting phenotypes; this switch affects the interaction between immune and host tissue cells to either drive fibrosis or mediate its resolution ^82^. Strikingly, *Lair1* (Leukocyte-Associated Immunoglobulin-like Receptor 1) was the only gene in our study, whose expression in two distinct organs—liver and brain—was linked to cognitive performance, potentially indicating a shared immune-regulatory thread along the liver–brain axis. LAIR1 is an inhibitory collagen/C1q receptor that tempers immune-cell activation; it directly restrains pro-fibrotic macrophage activity and promotes resolution of inflammation ^57,71^. Carroll et al.^83^ have shown that in neuropsychiatric systemic lupus erythematosus, an autoimmune disease that often causes cognitive impairments, microglial LAIR1 has been shown to be essential for dampening complement-driven synaptic loss, and down-regulation of microglial *Lair1* aggravates cognitive impairment. This study positions LAIR1 as a nodal checkpoint that decides whether microglial complement activity resolves or spirals into chronic synaptic loss ^83^.

Our results suggest that developmental processes may be triggered in concert with inflammation to compensate for compromised tissue homeostasis. The expression of MHC class Ib genes in the colon may reflect localized immune responses relevant to the immune pathway of the gut brain axis. Methylation and expression of several innate and adaptive immune genes strengthen this hypothesis. Taken together, these results indicate a clear involvement of immune and tissue repair and development processes in cognitive function. This observation is consistent with the chronic low-grade inflammation, often referred to as “inflammaging”, that accompanies age-related cognitive decline^84^.

At the metabolomics level, our main results indicate a role of taurinated bile acids in the maintenance of cognitive function. Bile acids, beyond their well-known role in digestion, have important signaling effects, influencing metabolism in the liver, gut, and other organs, including the brain ^85,86^. In the gastrointestinal tract, they interact with the gut microbiota and with host’s receptors to regulate metabolic and immune homeostasis ^87^. Bile acids have also been implicated in neurodegenerative diseases like hepatic encephalopathy and Alzheimer’s Disease (AD), with studies showing altered bile acid levels in AD patients compared to controls ^88–90^. *Glp-1* expression, suggested to be involved in the cognitive improvements seen in metformin-treated Type-2-Diabetes (T2D) patients ^91^, has been shown to be induced by taurinated deoxycholic acid (TDCA) in mice ^92^. It is therefore reasonable that the link between bile acids and cognition that we observed in our study could at least partially depend on the induction of *Glp-1*. Furthermore, Brunner’s glands, which are GLP-1 receptor-positive, play a significant role in the gut-brain axis by stimulating ileal mucin production ^93^. Notably, semaglutide, a GLP-1 receptor agonist used to treat T2D, has been associated with a reduced risk of a first-time Alzheimer’s diagnosis in a large cohort of T2D patients ^94^. An increased mucin production could potentially protect the intestine from colonization by harmful bacteria, decreasing gut permeability and providing an ecological niche for beneficial bacterial species. Studies in mice suggest that TDCA and its derivative tauroursodeoxycholic acid (TUDCA) can protect cognitive function in conditions such as Alzheimer’s disease and inflammation-induced memory impairment. For instance, TUDCA treatment has been shown to reduce amyloid-β deposition and preserve learning and memory in a transgenic Alzheimer’s mouse model ^95^. Similarly, in an LPS-induced neuroinflammatory mouse model, TUDCA lowered pro-inflammatory signaling and helped to maintain performance on memory tests ^96^. The findings reported in these studies point to a potential role of TDCA/TUDCA in preserving cognitive function. Despite our results not providing any direct link between bacterial metabolism and bile acid levels, it is well known that the gut microbiota can alter the bile acids pool, especially in terms of amination/deamination ^97^. While taurine-conjugated bile acids dominate the murine, but not the human, bile acid pool ^97,98^, the broader mechanisms of bile acid conjugation, deconjugation, and signaling are conserved and relevant in humans as well. As an example, bile acids conjugated with amino acids such as taurine or glycine can be detoxified more efficiently by making them more water soluble and therefore increase their excretion through urine ^97^.

By inferring active metabolic reactions within each mouse microbiome, we identified distinct cross-feeding of molecules —including heme, S-adenosyl-homocysteine, S-adenosyl-methionine, and D-ribose—as important features predictive of cognitive rank. Notably, none of the selected exchange reactions involved the gut lumen, suggesting that microbe–microbe interactions may dominate microbe–environment interactions in this context. Interestingly, most of these reactions are found in Bacteroidia (Fig.4D), suggesting that members of this class may play a central role in mediating community-level metabolic interactions associated with cognitive function. Bacteroidia are known to shape gut microbial community structure through the production of key metabolites that support other taxa ^99^. Consequently, our finding that cross-feeding reactions predominantly mediated by Bacteroidia emerged as key cognitive predictors underscores the central role of these bacteria in shaping the host’s metabolic environment—and, in turn, influencing cognitive outcomes during aging. Our results highlight the complexity of gut microbial networks and suggest that targeting key bacteria and their cross-feeding networks may offer new ways to support cognitive health.

On the level of each bacterium’s individual metabolic function, we observed a negative correlation between the estimated abundances of butyrate-producing reactions and cognitive functions in our aging mice cohort. This result suggests that the relationship between SCFAs and cognitive function is highly context-dependent, potentially influenced by factors such as age, microbiota composition, and inflammatory status. While SCFAs, including butyrate, are often considered beneficial for their anti-inflammatory and neuroprotective effects ^100,101^, findings by Sampson et al. ^41^ demonstrate that SCFAs can exacerbate neuroinflammation, microglial activation, and α-synuclein aggregation, leading to motor deficits in an α-synuclein-overexpressing mouse model of Parkinson’s disease. These results suggest that SCFAs can have negative effects in certain disease contexts. Other notable cognition-related microbial functions included methylglyoxal and folate metabolism, the former being implicated in neurodegeneration in the elderly ^44^, recently linked to damage in the blood brain barrier ^102,103^, and the latter being a vitamine whose role in cognitive maintenance during older age have been documented in both healthy subjects and Parkinson’s patients ^45,104^.

While microbial abundances and metabolic processes, as well as taurinated bile acids and 3-HK quantities, were identified as potential cognitive predictors in single-layer analyses, the multi-layer analysis indicated a stronger association between host features and cognitive function. Specifically, the integrative Boruta analysis retained 227 features (189 of them selected ten or more times) out of the 734 features identified at least once in the single-layer runs, and notably, all retained features originated from host tissues. A likely explanation for this reduction is that, unlike in the single-layer analyses, the candidate feature pool was pre-filtered for relevance, thereby reducing random noise. Additionally, because feature importance in random forests depends both on individual feature effects and on interactions with other variables, the integrated dataset likely altered the importance landscape through new inter-layer interactions that could not be evaluated in the single-layer analyses. As a result, only features with lower dependence on interactions with variables discarded in the single-layer runs, or participating in newly relevant cross-layer interactions, were retained. While this reasoning remains interpretative and was not formally tested here, it offers a plausible explanation for the observed reduction in feature number. Nevertheless, the observation that only host tissue features were ultimately retained suggests that the effects of the microbiome on cognitive function may be mediated via impacts on host tissue integrity and immune status, rather than directly through structural and metabolic differences between gut microbial communities. Examples of such indirect mechanisms include alterations in gut barrier permeability, which can enable the translocation of live bacteria or bacterial products into the bloodstream, contributing to systemic inflammation ^105^, as well as microbially-induced influences on the host’s immune status ^106^.

Our study also has some limitations. For experimental reasons, we included only male mice; future studies should incorporate both sexes to account for potential sex-dependent effects. While our Barnes Maze-based cognitive rank—being a robust and multifaceted indicator that combines spatial navigation, short- and long-term memory, as well as cognitive flexibility—provided valuable information about the cognitive status of each mouse, the requirement to collect tissues post-mortem for sequencing forced us to adopt a cross-sectional design. A longitudinal approach, tracking microbiome composition, fecal metabolomics, and cognitive performance across the lifespan, could provide critical insights into the dynamics of these parameters during aging. These trajectories could then be correlated with inflammatory and developmental signatures obtained at relevant endpoints, offering a more integrated view of cognitive aging and its microbial and systemic determinants. Finally, due to the inherent correlation between aging and cognitive decline, it is not possible to conclusively determine whether our findings are driven by age-related changes, cognitive function, or a combination of both. Although adjusting for age in our analysis was theoretically possible, age is the primary determinant of cognitive function in our cohort. Correcting for it would therefore eliminate much of the meaningful variation in cognitive performance, limiting our ability to identify relevant associations.

In summary, we have identified a comprehensive array of cognitive aging markers across diverse omics layers, incorporating both primary and model-derived data. Our findings identify key determinants of cognitive aging, underscoring the contribution of developmental and inflammatory processes, and adding further support to the concept that cognitive aging involves coordinated alterations in both central and peripheral systems, including the gastrointestinal tract. Furthermore, we present a rich dataset for further exploration of molecular contributors to cognitive decline, facilitating the development of targeted interventions against neurodegenerative processes.

## Methods

### 1. Animals and organ/feces collection

Male C57BL/6J/Ukj mice were maintained at 22°C on a 14 h/10 h day-night cycle and at a relative humidity of 55 +-10%. Mice were provided ad libitum access to ssniff mouse V1534-300 diet (ssniff Spezialdiäten GmbH, Soest, Germany) and drinking water. Mice were divided into five age groups: 3 (n = 16), 9 (n = 16), 15 (n = 16), 24 (n = 17), and 28 (n = 18) months old. The C57BL/6J/Ukj substrain lacks two common mutations found in the C57BL/6J strain: the DIP686 mutation in the crumbs family member 1 (*Crb1*) gene, and a mutation in the nicotinamide nucleotide transhydrogenase (*Nnt*) gene. Preserving these genes is relevant for metabolic and aging research, as the *Crb1* mutation impacts retinal integrity, while the *Nnt* mutation impairs mitochondrial NAD(P) transhydrogenase activity, potentially influencing oxidative stress responses.

We collected RNA sequencing data for brain, liver, and colon; methylation data through RRBS sequencing for brain and colon; shotgun metagenomics sequencing and untargeted metabolomics data of fecal content.

### 2. RNA seq

For the extraction of genomic DNA and RNA, the mice were sacrificed by cervical dislocation. Total RNA was extracted from tissue samples of the liver, colon, and left hippocampus using the Qiazol-chloroform extraction method with 1 mL of Qiazol Lysis Reagent (Qiagen, Hilden, Germany), as previously described in ^12^. RNA quality was determined via an Agilent Bioanalyzer 2100 using the RNA 6000 nano kit (Agilent Technologies, Santa Clara, CA, USA). An RNA Integrity Number (RIN) value of greater than seven was considered to be the minimum quality for sequencing. RNA libraries were prepared using TruSeq RNA stranded kit (Illumina, San Diego, CA) with polyA enrichment according to the manufacturer’s instructions. All libraries were sequenced on an NovaSeq 6000 machine (Illumina, San Diego, CA, USA) with an average of 36 million paired-end reads (2×100 bp) at Competence Centre for Genomic Analysis (CCGA, Kiel, Germany).

Illumina TruSeq adapter sequences were trimmed from forward and reverse reads using Cutadapt (2.8) with minimum sequence overlap of 3 bp, at most 10% mismatches allowed and minimum read length filter for 20 bp, as well as a two-color-chemistry aware, 3’-end quality trimming for a phred-score >= 25 (nextseq-trim=25). An additional quality filter was applied with Prinseq lite (0.20.4) for at most eight unknown nucleotides (‘N’) per read, an overall mean read quality of at least phred-score 15, and a minimum quality trim from both end of the read for a minimum phred-score of 12. Filtered reads were then mapped against *Mus musculus* reference genome (GRCm38.p6, mm10) via Hisat2 (2.1.0) with RNA strandedness set to FR, employing non-deterministic random seeds and suppressing the mixed alignments of read pairs. Only primary alignments for each read were kept, via a filtering step (-F 256) with Samtools (1.9). Gene counts were extracted using the tool featureCounts from the subread package (2.0.1) with reverse-stranded information, while ignoring chimeric fragments and allowing only properly mapped read pairs with insert sizes in between 50 bp to 600 bp. The colon RNA-Seq data generated in this study have been deposited in the GEO database under accession code GSE248002. The liver and brain RNA-Seq data generated in this study have been submitted to GEO and will be available under the accession code GSE302188.

### 3. RRBS seq

As in a previously published mouse aging study ^107^, we performed reduced representation bisulfite sequencing (RRBS) of 83 genomic DNA samples, from the mice’s colons and left hippocampi. The RRBS data generated in this study have been deposited in the Gene Expression Omnibus (GEO) database under accession code GSE233734. Genomic DNA was isolated with the DNeasy Blood & Tissue Kit (Qiagen). 200 ng of DNA was fragmented via a 5-hour Msp1-digest (New England Biolabs), followed by TrueSeq adapter ligation (Illumina, San Diego, CA, USA) and subsequent bisulfite conversion following the manufacturer’s protocol (EZ DNA Methylation Gold Kit, Zymo Research). The bisulfite converted DNA was amplified in a 19-cycles PCR with Pfu Turbo Cx Hotstart DNA Polymerase (Agilent Technologies), and the resulting libraries were quality checked using 4200 TapeStation (Agilent Technologies).

Read quality trimming (cutoff 20) and TruSeq universal adapter clipping was performed using Cutadapt v2.10. Due to MspI cutting site end-repair, R1 adapter was prefixed by NN and in case of paired-end data, unconditional 5’ clipping of length 2 was conducted on R2.

Mappings were done via segemehl v0.3.4 (methyl-C mode, accuracy 95%, reference GRCm38) and filtered for unique alignments. Paired-end alignments were further filtered for proper-read pairs, and overlaps of R2 with R1 were trimmed using BamUtil clipOverlap v1.0.14. Methylation rates were inferred utilizing haarz v0.3.0. Rates and coverages within a CpG context were compiled using BEDTools v2.30.0.

For the following analysis we calculated the mean promoter methylation values as the average methylation ratio of all CpG sites within a region of 1000bp downstream and 1000bp upstream of known transcription start sites in the mouse genome (GRCm38). The promoter methylation values of genes that were not covered in all samples had to be discarded due to the prerequisite of Boruta not allowing any missing values.

For plotting in Figures 5 and 6 we summarised promoter methylation on the gene level.

### 4. Non-targeted metabolomics using HILIC UHPLC-MS/MS

Fecal pellets were extracted with 1 mL of chilled methanol (−20 °C; LiChrosolv, Supelco; Merck KGaA, Darmstadt, Germany), homogenized and extracted with a Precellys Evolution Homogenizer (Bertin Corp., Rockville, MD, USA; 4,500 rpm, three 40-second cycles with a two-second pause). The samples were then centrifuged for ten minutes at 21,000 ×g and 4°C, and the supernatant was transferred into sterile tubes until analysis. Next, 100 μL of fecal methanolic extract was evaporated at 40°C with a SpeedVac concentrator (Savant SPD121P; Thermo Fisher Scientific, Waltham, MA, USA) and reconstituted in 75% acetonitrile (ACN; LiChrosolv, hypergrade for LC–MS; Merck KGaA) spiked with L-Leucine-5,5,5-d3 at 5 mg/L (99 atom % D; Merck KGaA), which was prepared in a methanol and water solution (50:50).

The samples were analyzed by using a UHPLC system (Acquity; Waters, Eschborn, Germany) coupled to a quadrupole time-of-flight (TOF) mass spectrometer (maXis; Bruker Daltonics, Bremen, Germany), as described previously ^108^.

Mass spectra were acquired at electrospray ionization of positive and negative modes (+/–). Spectrometric data were acquired in line and profile mode with an acquisition rate of 5 Hz from 50 to 1500 Da. Fragmentation experiments were set to the data-dependent mode (MS/MS [Auto]), where the three most intense ions were fragmented within one scan when a count reached over 2000. Ions were excluded after acquiring three MS/MS and reconsidered for fragmentation after six seconds. The collision energy was set to 20 eV for both modes with an isolation width of 8 Da. The electrospray ionization source parameters were as follows: capillary voltage of 4500 V for (+) and 4000 V for (–), end plate offset of (+/–) 500V, nebulizer gas of 2 bar, dry gas of 10 L/min, and dry heater of 200°C. Before measurements, the MS was calibrated using the ESI-L Low Concentration Tuning Mix (Agilent, Santa Clara, CA, USA). The ESI-L Low Concentration Tuning Mix (diluted 1:4 [v/v] with 75% ACN) was injected in the first 0.3 min of each run by a switching valve for internal recalibration by post-processing software.

Ammonium acetate (NH4Ac; LiChropur eluent additive for LC–MS; Merck KGaA) at 0.5 mol/L was adjusted to pH 4.6 with glacial acetic acid (Honeywell; Fluka, Seelze, Germany). Milli-Q water was obtained from a Milli-Q Integral Water Purification System (Billerica, MA, USA). Polar metabolites were separated by HILIC by using an iHILIC-Fusion UHPLC column SS (100 × 2.1 mm, 1.8 μm, 100 Å; HILICON AB, Umea, Sweden). The eluent compositions were as follows: Eluent A consisted of 5 mmol/L NH4Ac (pH 4.6) in 95% ACN (pH 4.6), and eluent B consisted of 25 mmol/L NH4Ac (pH 4.6) in 30% ACN. We started with 0.1% B, keeping it constant for two minutes, then increased B to 99.9% over 7.5 minutes. The condition of 99.9% B was kept for two minutes and reversed to 0.1% B within 0.1 minutes, held for 0.1 minutes. The run was completed after 12.1 minutes, and the column was equilibrated for five minutes before the next injection. The flow rate was set to 0.5 mL/min, the column temperature to 40°C, and the sample manager was cooled to 4°C; 5 μL of the sample was injected into the column (partial loop). The weak and strong washes consisted of 95% and 10% ACN, respectively.

### 5. Metabolite identification and metabolomic data processing

The raw LC–MS data were post-processed in GeneData Expressionist Refiner MS (version 13.5.4; GeneData GmbH, Basel, Switzerland), including chemical noise subtraction, internal calibration, chromatographic peak picking, chromatogram isotope clustering, valid feature filter (cut-off of 100 [+] or 2000 [−] maximum intensity and presence of features in at least 20% of samples for [+/−]), retention time range restriction (0.4–10.7 minutes), annotation of known peaks (mass-to-charge tolerance of 0.01–0.005 Da and retention time tolerance of 0.1), and MS/MS consolidation and export to merged MASCOT generic files (MGFs). Data processing resulted in a matrix containing features with mass-to-charge ratios (m/z), retention times, and observed maximum intensities for each sample. The data were normalized to the weighed-in wet fecal weight and a maximum intensity of 5 mg/L L-leucine-5,5,5-d3. The merged MGF files were used to search a spectral library using MSPepSearch (0.01 Da mass tolerance for precursor and fragment searches). Experimental and in silico spectral libraries were downloaded from MassBank of North America (https://mona.fiehnlab.ucdavis.edu/). Identification was performed by matching experimental MS/MS spectra against MS/MS of spectral libraries downloaded from MassBank of North America using MS PepSearch (release: 02/22/2019; 0.01 Da mass tolerance for precursor and fragment searches). From the MS PepSearch output, features with the highest dot product of the same identifier were retained, and then metabolites with a dot product of <500 were removed. Furthermore, Global Natural Products Social (GNPS) ^109^ Molecular Networking and Library Search were used to identify metabolites in the experimental MS/MS data. Therefore, for each feature from the metabolite data table, an MS/MS was selected based on the highest total ion count. The MS/MS with the highest count was submitted to GNPS (201 features for [+]. and 361 features for [−]). The GNPS Library Search was conducted with the following settings: The precursor ion mass tolerance and fragment ion mass tolerance were set to 0.01 Da, minimum matched peaks were set to 1, and the score threshold was set to 0.5. GNPS Molecular Networking was performed with the following settings: the precursor ion mass tolerance was set to 0.01 Da, the MS/MS fragment ion tolerance was set to 0.01 Da, and the cosine score was set to >0.5 with a minimum matched peak of 1, TopK was set to 10, the maximum component size was set to 100, the maximum shift was set to 200 Da, the minimum cluster size was set to 1, and the maximum analog search mass difference was set to 500. Metabolomics data for HILIC (+/-) have been made available at the MassIVE database (massive.ucsd.edu) with identifiers MSV000094409, and MSV000094410.

### 6. Cognitive Tests

Cognitive function in aging mice was assessed using the Barnes Maze test as in ^14^. Briefly, mice were habituated to the environment beforehand to minimize stress. The setup consisted of a 90 cm diameter platform elevated 100 cm above the floor with 20 holes, one of which led to an escape box. Bright light (1200 lux center, ≥900 lux periphery) served as an aversive stimulus. Following one habituation trial, mice underwent the following training protocol: three trials per day with 60-minute intervals over six consecutive days, using spatial cues to promote spatial navigation. Each trial lasted up to 240 s; if the escape hole was not found, mice were guided to it. On day 7, a probe trial was conducted without the escape box, followed four days later by a retention test with the escape box restored to its original location. Each of these consisted of a single trial allowing up to 240 s.

All experiments were recorded and maximum velocity was automatically calculated by the video tracking system EthoVision®XT 6.1 software (Noldus, Wageningen, Netherlands). Recorded parameters included primary latency, primary errors, velocity (mean and maximum), total distance, and cognitive score. Mice navigation strategy in the Barnes Maze was classified using the BUNS method (Illouz et al. ^110^), based on x-y tracking data from EthoVision. To avoid bias from post-escape exploration, only data up to the primary latency were analyzed, except in the probe trial. Each training trial and test was assigned to 1 of 6, and for the probe trial to 1 of 3 possible search strategies, which are rated with a numerical value defined as the cognitive score: highly spatial to non-spatial strategy: direct = 1, corrected = 0.75, long correction = 0.5, focused search = 0.5, serial = 0.25, and random = 0; probe trial highly spatial to non-spatial: focused search = 1, serial = 0.5, serial-random = 0.167. For mice who did not find the target hole during the probe trial, the cognitive score was set to 0.

### 7. Cognitive Rank

In order to assess the overall cognition of our mice we combined several readouts from the Barnes maze test procedure into a single score per mouse. Our final cognitive rank is comparable between mice within our study cohort, where lower ranks indicate worse performance in the Barnes maze and a generally lower cognition.

We decided to combine primary latency and cognitive score of the training period (days one to six), the probe trial on day seven and the retention tests on day eleven and twelve in order to reduce the impact of missing values and random noise. Due to the combination of the measures we can no longer distinguish between learning and memory retention, however we can still draw conclusions of the average cognitive ability of each mouse. In total, eight measures were combined, four from the training period, representing learning ability, and four from the following test trials, representing memory retention ability. Each of the eight measures was ranked individually within the entire cohort to obtain integer values from 1 to 83, where higher ranks represented better cognitive performance. An average rank was used for all tied values, and missing values received minimum rank. The median was taken across all eight measures for each mouse to obtain the final cognitive rank.

For the training period of six days with three trials per day we calculated a linear regression of the cognitive scores (see Methods “5. Cognitive Tests”) and used the slope and the y-axis intercept as one parameter each. Similarly, for the primary latency data of the training period, we used the slope and y-axis intercept of a logarithmic regression, accounting for the censored nature of the data, as tests were aborted if the mouse didn’t reach the escape hole after four minutes. This type of regression analysis was implemented in R using the VGAM package (version 1.1-12) with a vector generalized linear model and a Tobit regression model using -log(240 seconds) as lower and -log(one second) as upper limits. The code for this analysis is provided in the github. Further, from the probe trial at day seven we used the cognitive score and the primary latency as is. Finally the last two measures were obtained from the two retention test days eleven and twelve. For the primary latency we used the best (minimum) of both days for each mouse while for the cognitive score only data from day eleven was used.

### 8. Metagenomic sequencing

Microbial DNA was extracted from colon contents with the DNeasy PowerSoil Kit (Qiagen, Hilden, Germany) following the manufacturer’s protocol. Next, the DNA was prepared at the Max Planck Institute for Evolutionary Biology (Plön, Germany) with the Illumina NexteraXT Library Kit. All 83 samples were pooled and sequenced for 2 × 150 cycles in paired-end mode on all four lanes of an Illumina NextSeq 500 machine. Demultiplexing was performed with one mismatch allowed in barcodes. The raw read data were merged sample-wise and subjected to quality control for adaptor contamination and base call qualities. Adaptor sequences with an overlap of ≥3 bp and base calls with a phred+33 quality score of <30 were trimmed from the 3′ ends of reads using Cutadapt (version 1.12). Illumina’s Nextera transposon sequence and the reverse complement of TruSeq primer sequences were used as adaptor sequences. Subsequently, reads were subjected to quality control using Prinseq lite (version 0.20.4) with a sliding window approach that applied a step size of 5 bp, a window size of 10 bp, a mean base quality of <30, and a minimum-length filter that discarded any reads shorter than 50 bp after all other quality control steps. To filter out host sequences, the remaining sequences were mapped to the mouse reference genome (GRCm38.99) with Hisat2 (version 2.1.0). The remaining unmapped reads were then kept for MAG assembly.

Long-read sequencing was performed at the NGS core facility of the FLI Leibniz Institute on Aging (Jena, Germany) on colon contents of an independent mouse cohort from the same housing facility, consisting of 52 male mice of the same age and genotype, as previously described by Best et al. (2025)^12^. Briefly, DNA from age-matched samples was pooled, fragmented (75 kb) and size-selected for >6 kbp fragments. Each pool was loaded onto a SMRTcell and sequenced on a Pacific Biosystems RSII machine. The sequence output of this run had an average read length of 7.8–9.7 kb with a minimum yield of 750 kbp per pool/SMRTcell. The raw read data were subjected to quality control, processed into circular consensus sequences (CCS) and subreads (continuous long reads, CLR), and exported as FASTQ files via the SMRTportal (provided by Pacific Biosciences).

CCS can be considered to have better quality, since they are the base call consensus of at least three long reads. While CLRs usually have only one single polymerase read through, and thus lower base call confidence, CLRs can cover much longer sequence stretches. We therefore employed both CLRs and CCS to inform our downstream assembly. In order to keep only high quality long-reads we used the software Filtlong (version 0.2.0) which generates k-mer profiles from Illumina short read libraries of the same biological samples. Five million reads were randomly sampled from the corresponding illumina fastq-files via Seqtk (version 1.3-r114-dirty) for k-mer generation. Those k-mer profiles were then compared with those of our long-reads. Reads were split at regions of low k-mer similarity of at least 1,000 bp lengths (--split 1000). The 10 % of base pairs with the lowest quality were discarded (--keep_percent 90) and low quality 5’ and 3’ read ends were trimmed (--trim). Quality filtering preset “--length-weight” was used in order to put more emphasis on retaining longer reads versus reads with better k-mer identity. Finally a read length cutoff was employed to discard any CCS shorter than 1,000 bp and any CLR shorter than 10,000 bp. Following, we used the software MiniMap2 (version 2.20-r1061) with mapping profile “PacBio” to align the remaining CCS and CLRs against the mouse reference genome (GRCm38.99). With the software Samtools (version 1.10) we discarded all reads mapping to the mouse genome in order to retain only microbial reads and converted the unmapped reads back into fastq-format.

### 9. MAG assembly, binning and annotation

After quality control for low read quality, adapters, and host contamination, fastq-files were concatenated across all samples into a single large fastq-file for the Pacific Biosystems CCS and CLRs and into two large fastq-files for forward and reverse reads respectively for the Illumina shotgun reads. A full cohort assembly was done in OPERA-MS 0.9 using SPAdes 3.13.0 as short-read assembler, with k-mer sizes of 21, 33, 55, and 81 on the concatenated, quality-controlled read files. The assembly required a total runtime of ∼10,000 CPU-hours and resulted in a total assembly size of 2.3 Gbp across 335 thousand scaffolds (>= 1,000 bp) with a scaffold N50 of 17,336 bp and the scaffold length ranging from 1 kbp - 1.8 Mbp. Cutoffs for filtering assembled scaffolds were determined in R statistics software to be coverage ≥37 and length ≥1000 bp and used to filter out scaffolds via python (https://github.com/APDLS/CVLFilter) as described by Douglass et al. in ^111^. The quality-controlled metagenomic Illumina shotgun reads were mapped back to the filtered scaffolds with Hisat2 (version 2.1.0); the insert size was 0–1,000 bp in the non-deterministic, “fr” stranded mode with end-to-end and no spliced alignment. Non-unique mappings and unaligned reads were discarded. The scaffold coverage depth was determined with the jgi_summarize_bam_contig_depths script from MetaBAT (version 2.12.1). This coverage depth information was then used to sort the remaining scaffolds into bins, each representing single bacterial genomes, with the binning tools MetaBAT (version 2.12.1), Autometa (version 2.0a0), CONCOCT (version 1.1.0), and MaxBin (version 2.2.7). For CONCOCT, the scaffolds were broken up into 10 kbp chunks. Bin refinement was conducted with the combined results of all four binners (383 bins) with DASTool (version 1.1.2); subsequently, quality metrics were calculated by CheckM (version 1.1.2). 249 bins with a quality estimate of >=90% and a contamination estimate of <=5% were considered for further analysis and are henceforth referred to as MAGs. The MAGs were taxonomically annotated with GTDB-Tk (version 2.1.1) and database version r214. The complete characterization of the MAGs is provided in Supplementary Table 3. Metagenomic raw read and MAG assembly data were deposited in the European Nucleotide Archive (ENA) and will be made available under BioProject PRJEB90935.

### 10. Bacterial models reconstruction with gapseq

For each MAG, a COBRA model has been generated using gapseq version: *1.2 e3210c0f* and sequence DB md5sum: *bf8ba98* using a bitscore threshold of 150 for gene identification. The medium used for each model’s gapfilling is based on the SSNIFF molecular diet, that reflects the quantities of each molecule in the SSNIFF food pellets based on the vendor’s information and on the roadmap presented in ^112^, similarly to the diet that has been used in ref. ^12^.

### 11. FVA-based reaction abundances

Flux variability analysis (FVA) was employed to determine the possible flux ranges for each reaction in the metabolic models of each bacterium separately. Biomass production was defined as the objective function, with flux ranges calculated to support near-maximal growth (99% of the maximum growth was used as a threshold). A binary incidence matrix was then created, where columns represented bacteria and rows represented biochemical reactions. Reactions exhibiting non-zero fluxes (>10⁻⁶) were assigned a value of “1.” This matrix was normalized based on the number of active reactions per bacterium and subsequently weighted by the bacterial abundances in each mouse. The resulting reaction abundance matrix for each microbiome was normalized by the total reaction scores per mouse, enabling comparisons of relative reaction abundances across mice.

### 12. Bacterial community simulations

For each microbial community obtained from fecal samples, community metabolic models were reconstructed using a coupling approach (MicrobiomeGS2^18^, an R package). This was achieved by combining the gapseq models associated to each community into merged models, where the bacterial abundances served as coupling constraints. In community models, bacterial models have individual compartments, and all models are interconnected through an environmental compartment allowing for the flow and exchange of metabolites. Thereafter, a community-level biomass reaction was introduced that accounts for the individual bacterial biomasses according to their relative abundance in the respective sample. An objective function was set to optimize for the community growth. cplexAPI R package, which employs CPLEX solver with an academic license ^113,114^, was used to conduct the metabolic simulation.

### 13. Feature selection with Boruta

Multi-omics integration in our study involved a much higher number of variables compared to the number of data points, sparse yet meaningful signals, and strong correlations both within and across layers. Standard univariate testing would explode the hypothesis space and force prohibitively strict multiple-testing correction. To obtain a biologically informative feature set we therefore adopted Boruta, a random-forest wrapper designed to retain every variable that carries information rather than a minimal subset ^19^. Boruta is widely applied in gene expression and microbiome studies, where it has been shown to capture both linear and non-linear associations and to resist over-fitting on artificial data ^115^. In brief, Boruta analyzes the importance of each feature by executing multiple random forests, in which each individual random-forest run augments the dataset with “shadow” variables obtained by permuting the original columns. A feature is confirmed when its mean importance score is significantly higher than the best shadow feature. This comparison provides an implicit multiple-testing correction, so no additional FDR control is required ^19^.

Before applying Boruta, very low-variance variables were discarded from all data layers using the nearZeroVar function from the R package *caret*. After variance filtering, the host tissues dataset consisted of 31530, 30286 and 26307 gene expression features for the hippocampus, colon and liver respectively, and 54583 and 61370 features from the hippocampus and colon methylation layers. The metabolomics dataset contained 1962 variance-filtered features, while the microbial datasets contained 249, 167, and 2592 from the MAG abundances, community-modelling, and reaction abundance tables, respectively. Additionally, all samples with unavailable data in at least one data layer were excluded from subsequent analysis. A total of 74 samples were retained for all subsequent analyses including correlations and plots. Boruta was first applied separately to each omics data layer. Each omics layer was processed in 50 independent Boruta runs. Features confirmed in ten or more runs were deemed robust and used for single-layer biological interpretation. The union of every feature that appeared in at least one run served as the candidate pool for an integrated analysis; the obtained dataset was again subjected to 50 Boruta runs, and features confirmed in ten or more runs were retained as cross-layer predictors. Since Boruta evaluates feature relevance relative to randomly permuted shadow features generated from the input dataset, integrating multiple omics layers alters the shadow feature composition compared to single-layer analyses, potentially yielding a different selection stringency. Additionally, as feature importance in random forests depends on both the feature itself and its interactions with other variables, integrating multiple data layers modifies the context in which feature relevance is assessed. In our setting this resulted in 227 features being selected at least once during the analysis of the integrated dataset, from the 734 features identified across individual data layers.

Our objective is feature discovery rather than predictive benchmarking. Consequently, we did not perform an additional cross-validation step for model accuracy; all stability assessments derive from the repeated Boruta runs themselves.

### 14. Gene Ontology enrichment analysis

GO enrichment was conducted using the *enricher* function from the *clusterProfiler* R package, with a minimum gene set size of 10, maximum of 500, and Benjamini-Hochberg correction for multiple testing. The background set for the expression or methylation matrix of each tissue served as its universe. Enriched GO terms were ranked by adjusted q-value and GO category size, prioritizing more specific terms. Only GO terms with a nominal p-value ≤ 0.05 and at least three associated genes were retained.

No gene set was significantly enriched in cognition-associated genes from the colon transcriptomics data; in order to interpret this data, we manually checked which cognition-associated genes are known or predicted to be a transcription factor according to the Transcription Factor checkpoint 2.0, a database that incorporates information from twelve collections of transcription factors ^116–127^.

### 15. Retrieval of genetic functional information

In addition to the information obtained from the Transcription Factor checkpoint 2.0, genes and loci identified in this study were cross-referenced with multiple literature and genomic databases to ensure biological relevance, particularly focusing on neurodegeneration, inflammation, and gut-brain axis regulation. Primary sources included the Mouse Genome Informatics Database and the Mouse Phenotype Database ^128,129^,GeneCards ^130^, and IMGC ^131–134^ all of them offering detailed phenotype annotations. These resources allowed interpretation of gene functions relevant to the genes that we identified in our study.

### 16. Data preparation for Graphlan

A phylogenetic tree of the 249 high-quality MAGs was generated and visualized with GraPhlAn ^135^. We divided the relevant features obtained from the reaction abundances layer into seven manually defined functional groups based on the metabolites involved in each reaction (butyrate, rhamnose, propanediol, xylulose, uracil, methylglyoxal, and folate) and one group containing all Boruta-selected community modelling exchange reactions. We then calculated a score for each functional category for each MAG dividing the number of reactions that each metabolic model included, over the total number of reactions in the category. Functional categories that were equally represented in all models (propanediol) were excluded from the plot. For each MAG, median relative abundance across the cohort was calculated, and associations between abundance and cognitive rank were assessed using Spearman correlation. These features were used to annotate the phylogenetic tree with both functional and abundance metadata.

## Data and Code availability

Data and code used in this publication are available under the doi: 10.5281/zenodo.15849341

## Acknowledgements

We acknowledge funding by the German Research Foundation to C.K. within the scope of CRC1182 (project A1.5), the Research Group miTarget (FOR5042) and the Cluster of Excellence ‘Precision Medicine in Chronic Inflammation’ (EXC2167). We also acknowledge funding from the German Research Foundation (DFG, project number: 416 418087534) to C.K., C.F. and S.H., the Carl Zeiss Foundation IMPULS programme (project number: P2019-01-006) to C.F. and S.H., and the European Union’s Horizon 2020 Research and Innovation Programme (under the Marie Sklodowska-Curie grant: agreement number 859890 (SmartAge)) to C.K., C.F. and S.F. This work was supported by the DFG Research Infrastructure NGS_CC (project number: 407495230) as part of the Next Generation Sequencing Competence Network (project number: 423957469). Short-read sequencing was conducted at the Competence Centre for Genomic Analysis (Kiel, Germany). We thank K. Cloppenborg-Schmidt for the excellent technical assistance in preparing the samples for metagenomic sequencing. We thank Silke Szymczak for the valuable suggestions about the design of the feature selection procedure.

## Author contributions

S.F. calculated the reaction abundances, led the manuscript writing, literature review and interpretation of the results, and created the figures. T.D. implemented the Boruta-based feature selection, with initial support from S.F. for the conceptual design. L.B. coordinated the exchange of data and samples between participating laboratories, designed the cognitive rank metric, performed the gene ontology enrichment analysis, and, together with T.D., assembled the bacterial genomes. G.M. provided the dietary information for the metabolic models. J.Z. built the metabolic models. A.S.K. conducted the community modelling simulations. M.H., R.S., and S.E. conducted the Barnes Maze experiments, processed the data using the BUNS algorithm, and collected biological samples. J.T. provided extensive feedback on the manuscript. M.O., K.R., S.Fr. and S.H. carried out the RRBS sequencing and preprocessed the data. A.W. and P.S. performed the metabolomics analysis. S.Fr. performed RNA sequencing. J.B. conducted metagenomic sequencing. C.F. and C.K. conceived and supervised the study. All authors reviewed and approved the final version of the manuscript.

## Competing Interests

The authors declare no competing interests.

## Notes

https://www.ncbi.xyz/geo/query/acc.cgi?acc=GSE248002

https://www.ncbi.xyz/geo/query/acc.cgi?acc=GSE233734

doi:10.25345/C5F47H53G

doi:10.25345/C5JW86Z7W

doi:10.5281/zenodo.15849341

### Competing Interest Statement

The authors have declared no competing interest.

